# How Nasal Cavity Structure Influences Empty Nose Syndrome Severity: A CFD-Based Analysis

**DOI:** 10.1101/2025.04.04.645990

**Authors:** Aurelien Rumiano, Tuan Dinh

## Abstract

**Background:** Empty Nose Syndrome (ENS) is a debilitating condition that can occur after partial or total turbinectomy, leading to impaired nasal airflow sensation, breathing difficulties, and sleep disturbances. While ENS is often diagnosed using the ENS6Q questionnaire, its precise causes remain unclear. Some patients with significant turbinate loss develop minor ENS symptoms, whereas others experience severe symptoms after minor mucosal cauterization. Understanding the structural and aerodynamic factors contributing to ENS is crucial for improving diagnosis and prevention.

**Objective:** This study aims to identify correlations between the ENS6Q score and key anatomical and aerodynamic parameters obtained from computational fluid dynamics (CFD) simulations in ENS patients.

**Methods:** We reconstructed patient-specific nasal cavity models from computed tomography (CT) scans and performed CFD simulations. The analysis focused on five key parameters: the remaining turbinate volume, total mucosal surface area, nasal resistance, average cross-sectional area, and airflow imbalance between the two nasal cavities. These parameters were then compared to ENS6Q scores.

**Results:** Preliminary findings suggest that a lower remaining turbinate volume, reduced mucosal surface area are associated with higher ENS6Q scores. Additionally, significant airflow asymmetry between the two nasal cavities appears to correlate with more severe symptoms.

Furthermore, our data indicate that individuals with larger nasal cavities and greater preoperative mucosal surface area tend to be more resilient to turbinectomy. For an equivalent amount of turbinate resection, patients with initially smaller nasal cavities thus having less mucosal surface experience more severe ENS symptoms.

**Conclusion:** By quantifying the anatomical and aerodynamic characteristics of ENS patients, this study provides new insights into the structural factors contributing to ENS severity. These findings may help refine diagnostic criteria and guide surgical approaches to minimize ENS risk.

## Introduction

First described by Kern and Stenkvist (1), Empty Nose Syndrome (ENS) refers to a set of symptoms that can occur after partial or total turbinectomy.

The turbinates are made up of bony blades surrounded by erectile tissue and finally covered with mucous membrane. Their purpose is to humidify, filter, and heat the air before it reaches the lungs. They also inform the brain of the passage of airflow via TRPM8 receptors (2).

Partial or total removal of the turbinates can create empty nose syndrome; simple cauterization of the mucous membranes without removing volume can also cause it due to the alteration of the health of the mucosa.

People suffering from empty nose syndrome have problems feeling the passage of air, which causes a feeling of suffocation and disrupts breathing control and sleep (2). Nasal dryness, feeling of cold air, and infections are also common symptoms.

Recent advances in computational fluid dynamics (CFD) have enabled the study of these airflow patterns. CFD simulations can quantify critical parameters such as nasal resistance, airflow velocity, and wall shear stress (WSS), an indicator of mucosal stimulation. Several previous studies have used CFD to demonstrate that patients with ENS have lower nasal resistance and altered airflow distribution, as well as reduced stimulation along the inferior meatus. (4)

In this study, we perform CFD simulations on nasal cavity models based on computed tomography (CT) scans of patients with ENS to characterize their airflow patterns and compare these results with symptom severity. Our goal is to better understand the pathophysiology of ENS, with the aim of informing surgical strategies that could reduce the risk of developing ENS.

## Problem statement

To diagnose empty nose syndrome two questionnaires are often used ENS6Q and SNOT 20. From a certain score, the person is considered to suffer from ENS.

Some people who have had part of their turbinates removed develop milder symptoms than others with a similar extent of turbinate resection. Therefore, we still do not fully understand what determines the severity of symptoms in empty nose syndrome.

However, we already have a general understanding of what ENS causes. Two main points.

The more turbinate tissue is removed, the higher the risk of developing ENS.

The condition of the nasal mucosa also plays a role, the more tissue that remains but is non-functional, the higher the risk of developing symptoms.

## Aims

This study aims to shed light on the precise causes of ENS by finding correlations between the ENS6Q score and various data taken from fluid simulations of a cohort of patients whose turbinates have been partially or completely cut.

## Methodology

We simulated the passage of air through the nasal cavity of 19 people using a suite of open source software tools.

Each patient had to complete an ENS6Q questionnaire to assess their symptoms.

### Study Design and Patients

This prospective computational study analyzed CT scans from patients clinically diagnosed with ENS. Inclusion criteria were: 1/a history of partial or total inferior turbinate resection with persistent ENS symptoms for at least six months post-surgery; 2/availability of high-resolution CT imaging suitable for three-dimensional reconstruction; and 3/absence of other significant nasal pathology. ENS symptom severity was quantified using the validated Empty Nose Syndrome 6-Item Questionnaire (ENS6Q). When available, data from healthy control subjects (with no history of nasal surgery or chronic nasal complaints) were used for comparison.

### Nasal Airway Modeling

The patients CT scans were converted to 3D in STL format using the open-source software SLICER 5.2. Part of the air surrounding the nostrils and part of the pharynx were kept.

The meshing was performed using the OpenFoam 1812 software, with each model containing between 500,000 and 1 million cells depending on the case. Beyond 1 million cells, we did not notice any differences in results during the simulations.

### CFD simulation

CFD simulations were conducted with a finite-volume solver (OpenFoam, 1812), assuming steady-state, incompressible airflow under inspiratory conditions at a flow rate of 15 L/min. Air was modeled as a Newtonian fluid (density ∼1.204 kg/m^3^, dynamic viscosity ∼1.8×10^−5^ Pa·s).

A flow rate inlet boundary condition was applied at the nostrils, and a constant pressure (0 Pa gauge) was set at the nasopharyngeal outlet.

Solver: SIMPLE

Turbulence model: RANS k epsilon.

The walls’ temperature and humidity parameters were not taken into account. Given velocities that do not exceed 5m/s, air is considered incompressible.

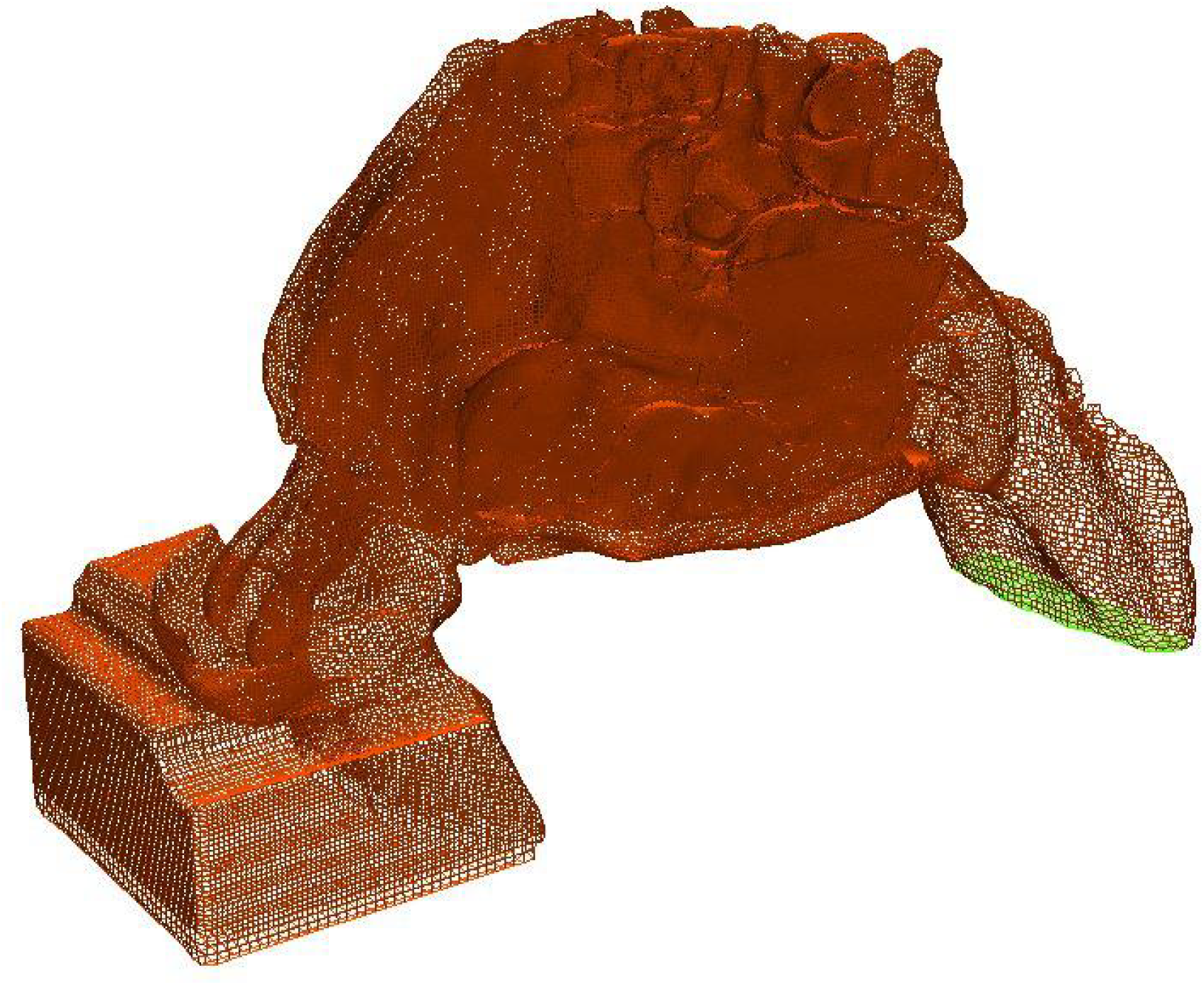

Example of meshing a nasal cavity

Finally, the data on nasal resistance, cross sections, etc., were retrieved with Paraview 13.0 software.

With the data, we created graphs to visualize possible correlations between the ENS6Q score and each piece of data. We chose the point cloud with a regression line.

Each patient received a detailed simulation report, including 3D views of Wall Shear Stress, airflow, and cross-sections of airflow velocity.

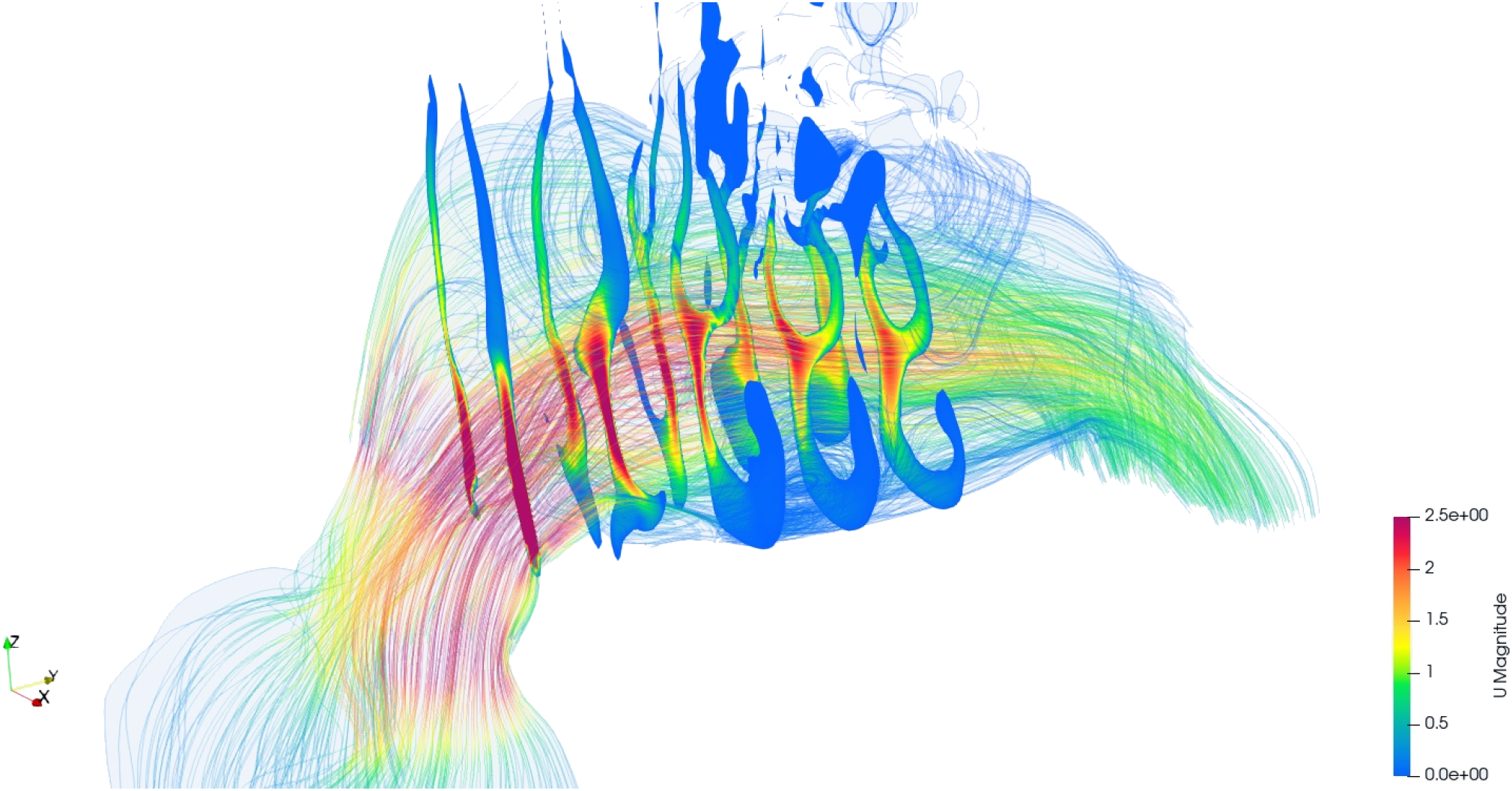

View of air flow with sections, scale up to 2.5 m/s

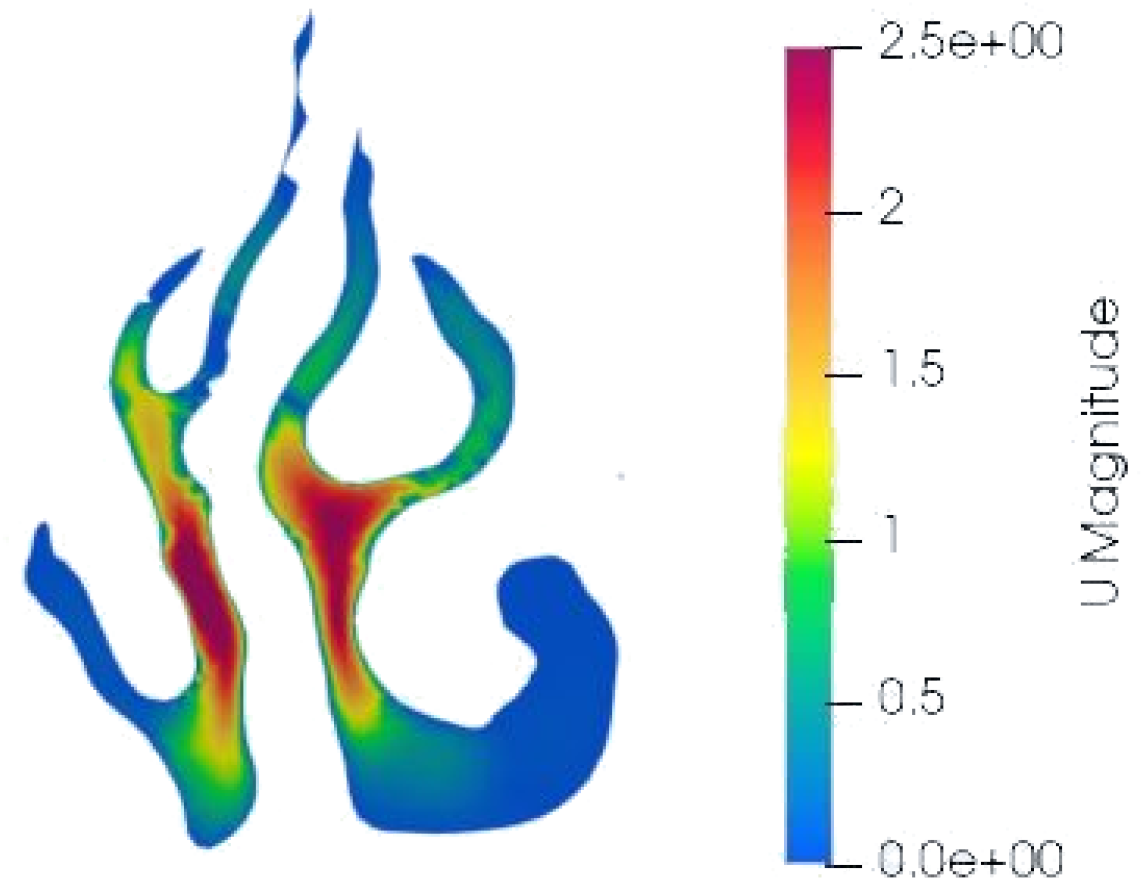

Air speed cut with scale up to 2.5 m/s

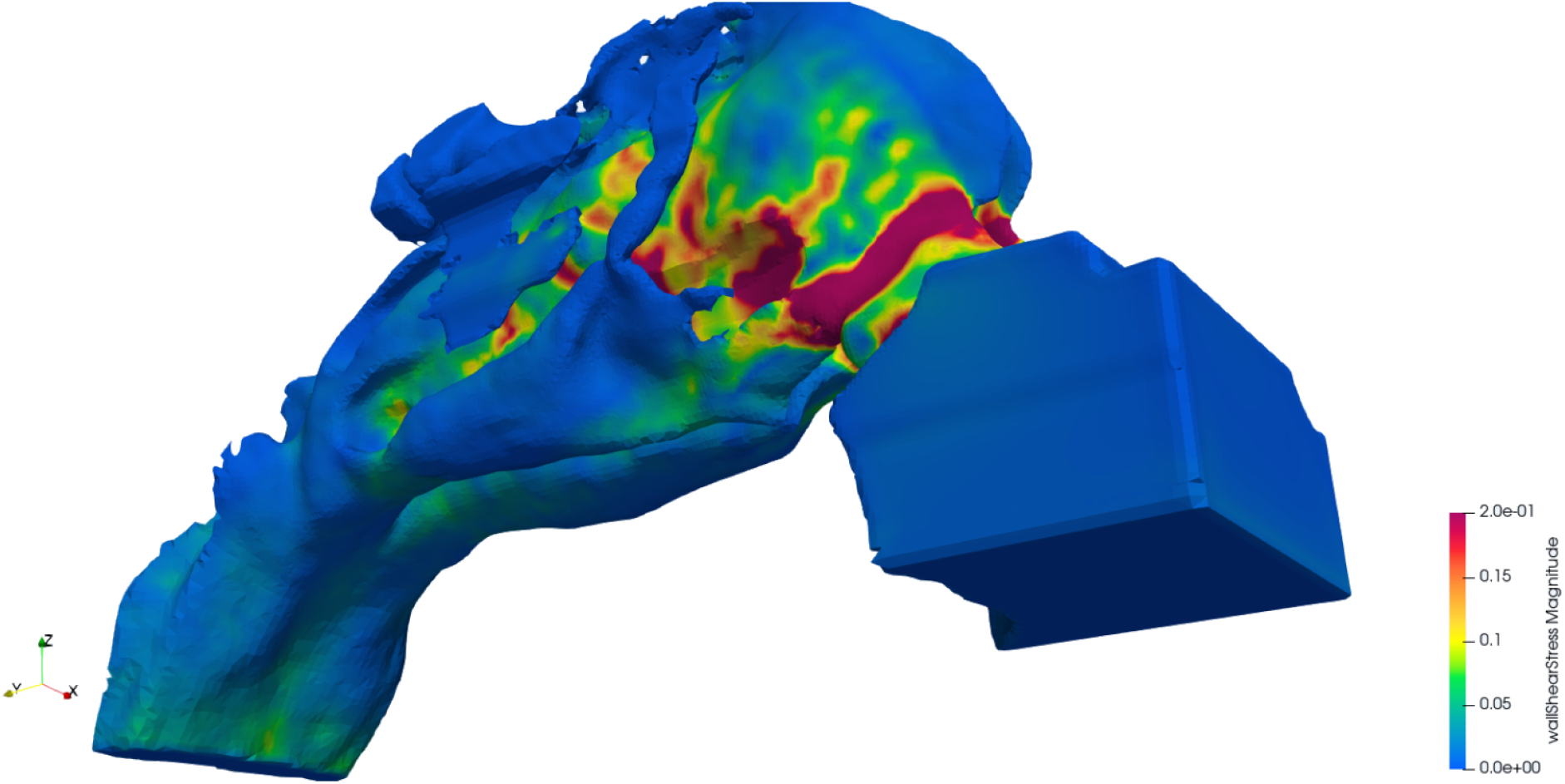

View of Wall Sheer Stress with a scale of up to 0.2 Pa

[Example of report sent to patients](https://enstips.com/aurelien/).

## Results

19 ENS patients (17 males, 2 females) were included. All patients had undergone partial or total inferior turbinate resection, with some having additional nasal surgeries. The median ENS6Q score was 16 (range: 6–24), indicating moderate to severe symptoms. Common complaints included a sensation of suffocation, nasal dryness, and crusting.

The fluid simulations allowed us to extract the data that seemed most relevant to us, namely: The total nasal resistance, the flow imbalance between the left and right nasal cavities, the average of the cross sectional area, the average perimeter of the quantity of mucosa which in fact reflects the surface of the total mucosa along the nasal cavity. We were obliged to take the perimeter since the measurement was carried out in sections. Finally, the percentage of remaining turbinates was quantified visually. We did not collect data on WSS because they are difficult to quantify; the peak WSS often used in studies on ENS is subject to error.

| Subjects | deltaP (Pa) | Airflow imbalance (ratio) | Average cross sectional area (mm<sup>2</sup>) | Percentage remaining turbine (left) | Percentage remaining turbines (right) | Average remaining turbine (%) | Average perimeter (mm) | ENS6 Q total |
| --- | --- | --- | --- | --- | --- | --- | --- | --- |
| 1 | 10.50 | 1.27 | 320 | 50 | 0 | 25 | 306 | 19 |
| 2 | 4.9 | 2.34 | 370 | 90 | 0 | 45 | 314 | 20 |
| 3 | 8 | 1.67 | 327 | 70 | 90 | 80 | 398 | 10 |
| 4 | 8.7 | 1.12 | 412 | 70 | 10 | 40 | 304 | 10 |
| 5 | 6.8 | 2.19 | 345 | 20 | 70 | 45 | 314 | 17 |
| 6 | 10.2 | 1.10 | 307 | 90 | 70 | 80 | 353 | 6 |
| 7 | 4.1 | 1.05 | 500 | 90 | 70 | 80 | 282 | 11 |
| 8 | 6.75 | 3.00 | 291 | 90 | 70 | 80 | 290 | 16 |
| 9 | 5 | 1.05 | - | 40 | 30 | 35 | - | 16 |
| 10 | 7.2 | 2.17 | 336 | 100 | 80 | 90 | 355 | 20 |
| 11 | 12.5 | 1.10 | 315 | 40 | 50 | 45 | 276 | 26 |
| 12 | 13.5 | 1.50 | 275 | 80 | 90 | 85 | 304 | 10 |
| 13 | 3.6 | 1.27 | 492 | 50 | 50 | 50 | 311 | 10 |
| 14 | 12.7 | 1.00 | 245 | 70 | 70 | 70 | 301 | 19 |
| 15 | 6 | 1.10 | 399 | 90 | 90 | 90 | 366 | 12 |
| 16 | 8.3 | 1.25 | 301 | 70 | 70 | 70 | 289 | 22 |
| 17 | 5 | 1.40 | 360 | 20 | 80 | 50 | 260 | 17 |
| 18 | 7.6 | 1.20 | 252 | 80 | 90 | 85 | 281 | 13 |
| 19 | 12.75 | 1.90 | 201 | 90 | 50 | 70 | 216 | 24 |

Full data pulled from the simulations

Full data pulled from the simulations
|  | r | p-value |
| --- | --- | --- |
| DeltaP | 0.268 | 0.267 |
| Airflow imbalance | 0.246 | 0.31 |
| Average cross sectional area | -0.417 | 0.0848 |
| Remaining turbines | -0.297 | 0.217 |
| Mucosal perimeter | -0.5 | 0.034 |

Among the analyzed parameters, mucosal perimeter shows the strongest negative correlation with ENS6Q (r=−0.5, p-value=0.034), The more mucosa remains, the lower the ENS6Q score is, suggesting that individuals with a greater initial mucosal surface may be more resilient to the effects of turbinectomy, experiencing fewer symptoms at the same level of procedural aggressiveness. The average cross-sectional area also shows a moderate negative correlation (r=−0.417), with a p-value of 0.0848, which is close to being statistically significant. Other parameters exhibit weaker correlations and do not reach statistical significance. These findings highlight the potential impact of mucosal structures on symptom severity in ENS.

### Pressure loss

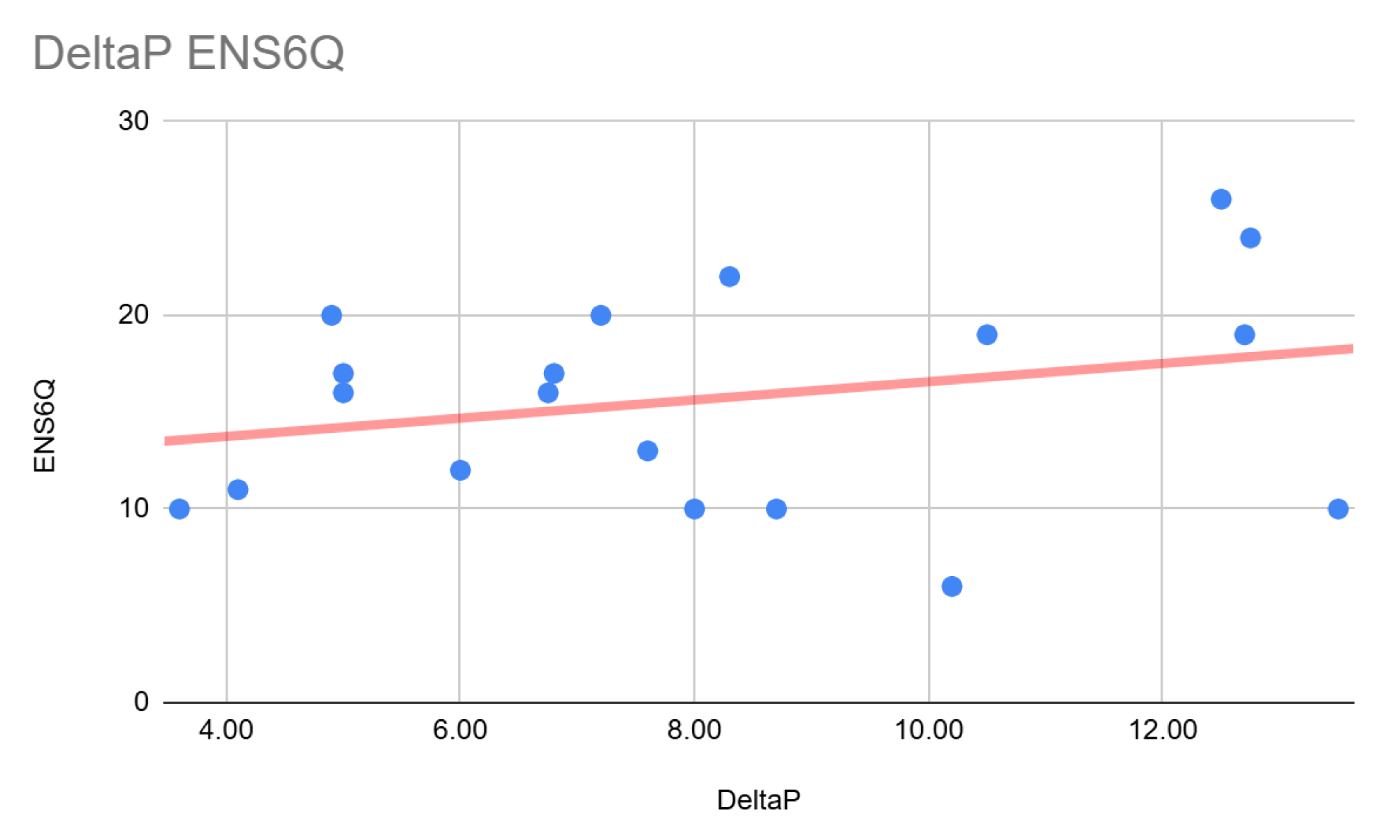

The DeltaP represents the pressure loss or load loss that the nasal cavity plus part of the pharynx causes during the inspiratory phase. The unit is in Pascale (Pa). It can be compared to nasal resistance.

This graph shows a weak correlation between nasal resistance and the ENS6Q score (r = 0.268), which is not statistically significant (p-value = 0.267). Interestingly, higher resistance is associated with a higher symptom score, which might seem counterintuitive at first. One might expect opposite results given that the reduction in volume leads to a reduction in nasal resistance. Several factors can explain this result.

The simulation included the pharynx, which modifies the result; for example, a small pharynx increases the total resistance. A more precise assessment of nasal resistance might have been obtained by excluding the pharynx and limiting the analysis to the choana.

### Quantity of turbinates remaining

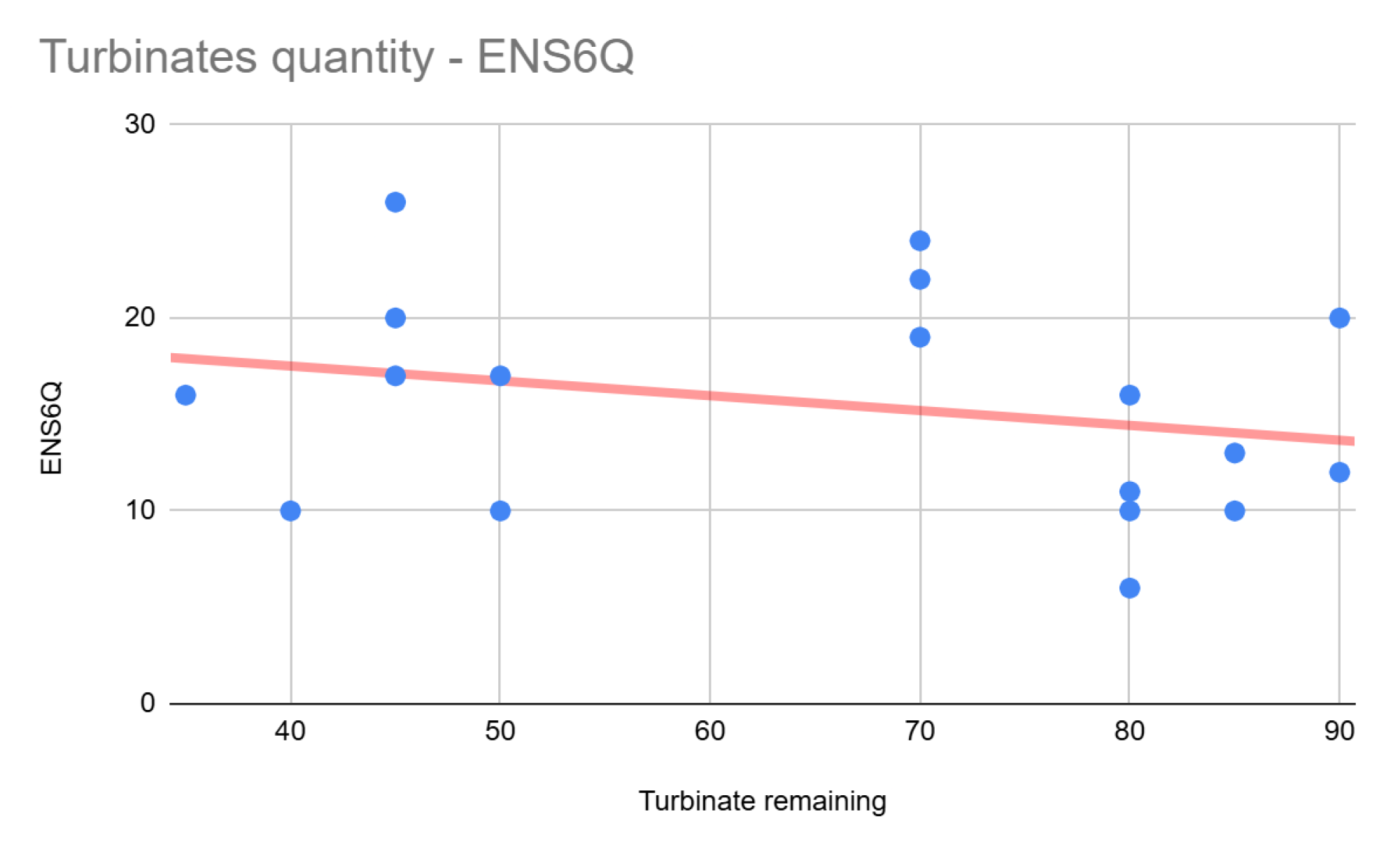

The quantity of remaining turbinates was quantified visually because no software can measure this data.

Here, we observe a weak negative correlation between the percentage of turbinates remaining and the ENS6Q score (r = −0.297, p-value = 0.217). Although this suggests that having more turbinates is associated with fewer symptoms, the correlation is weak and not statistically significant. These results are not surprising and reflect that the aggressiveness of turbinectomy affects the severity of symptoms, but further studies with larger sample sizes may be needed to confirm this trend.

### Mucosal surface

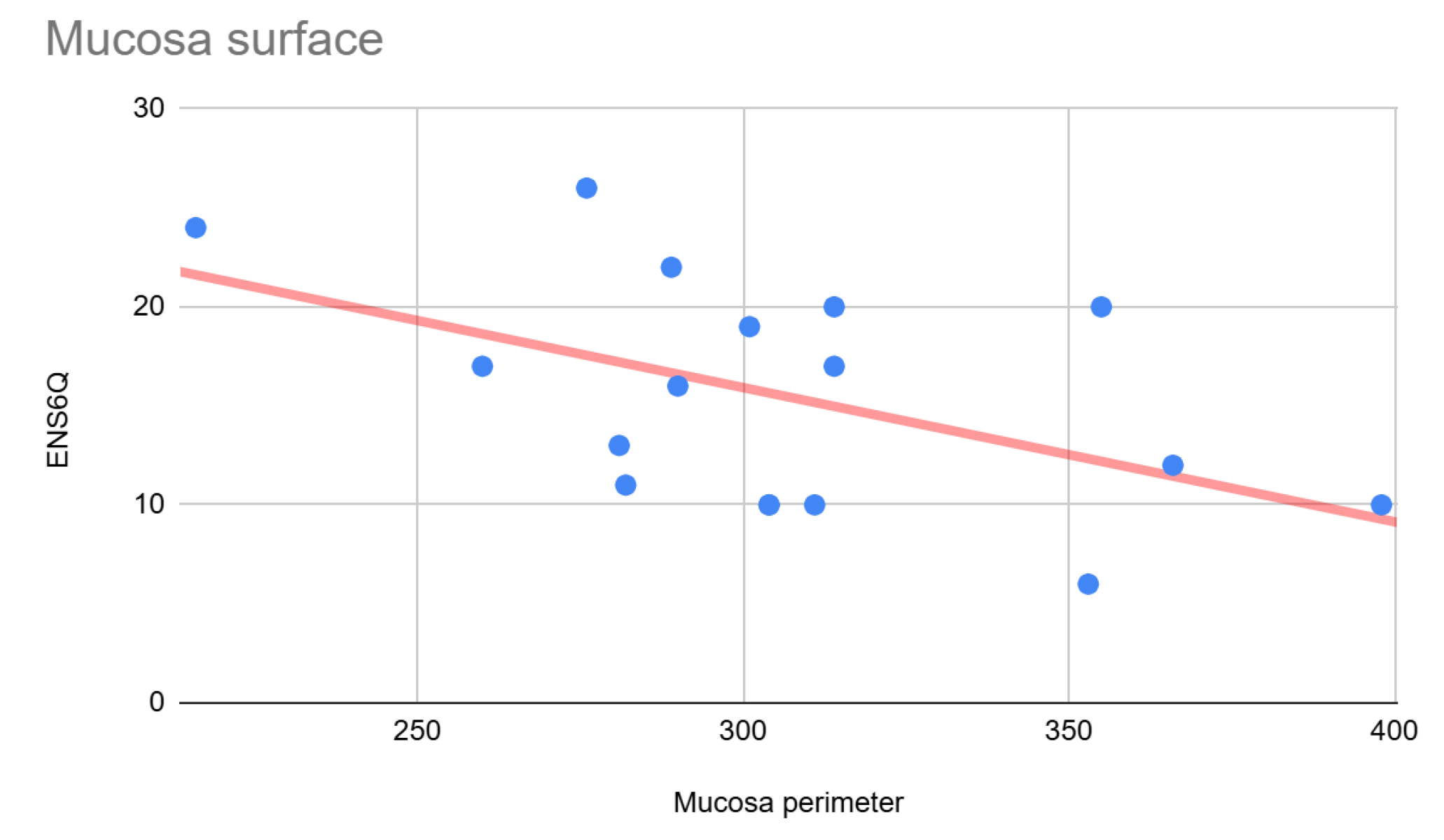

The quantity of mucous membrane was measured as follows:

We made three cuts 20 mm from the nostrils: one at 20 mm, one at 30 mm, and one at 40 mm. Then, we measured the perimeter and finally calculated the average. The result obtained is, therefore, an average perimeter in mm, representing the mucosa’s surface between 20 and 40 mm from the nostrils. We chose these cuts because they represent the center of the nasal cavity, where the turbinates are generally cut the most during turbinectomies.

Here, the correlation is even clearer, and we think it is the most relevant result of the study. The mucosal perimeter shows the strongest negative correlation with the ENS6Q score (r = −0.5, p-value = 0.034), and this result is statistically significant.

The more mucosa remains, the lower the ENS6Q score is, independent of all other results. That is to say, a person who has undergone a more aggressive turbinectomy than another but has a higher amount of remaining mucosa will still have lower symptoms. The variation in the quantity of mucosa in a nasal cavity is explained not only by the quantity of remaining turbinates but also by the size of the nasal cavity, which is largely correlated to the width of the dental arch and, therefore, of the palate.

This could, among other things, explain why some people experience fewer symptoms than others with the same amount of cut turbinates. Of course, this is one parameter that can explain this phenomenon, but it is not the only one.

### Average cross-sectional area

The section average was calculated as follows:

We measured the cross-sectional area at intervals of 10 mm, from the nostrils to a depth of 40 mm. The average of these measurements was then calculated, and the result is expressed in mm^2^.

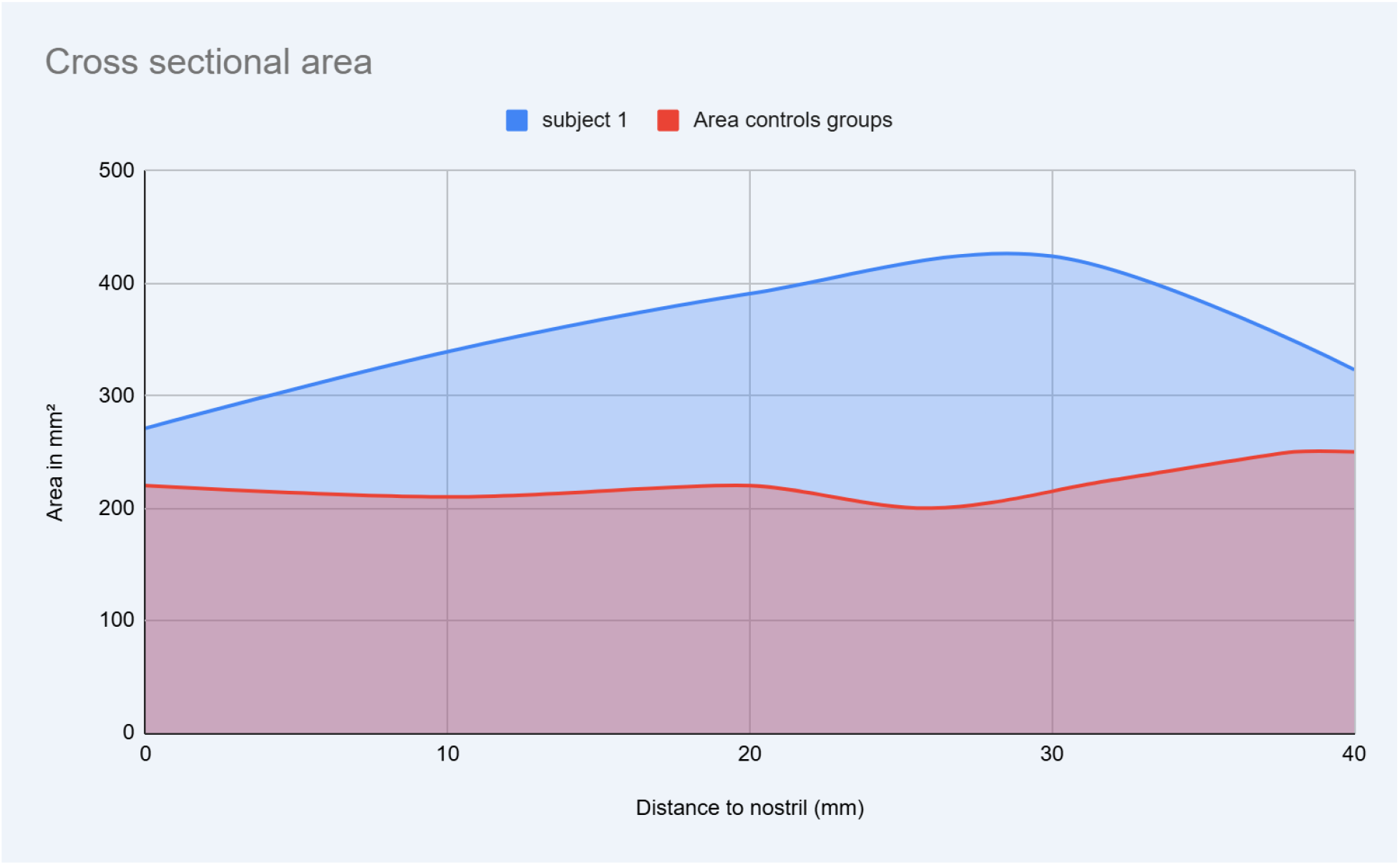

Example of cross section reading compared to a control group (3)

Surprisingly, we observe a negative correlation between the cross-sectional average and the ENS6Q score (r = −0.417, p-value = 0.0848), which is relatively strong and approaches statistical significance. This seems counterintuitive, given that turbinectomy increases cross-sectional area and reduces nasal resistance. However, this can be explained. Subjects with a larger/more developed nasal cavity also have a higher cross-section and, therefore, more mucosal surface area. As we saw previously, the amount of mucous membrane is correlated with the reduction of symptoms.

### Airflow imbalance

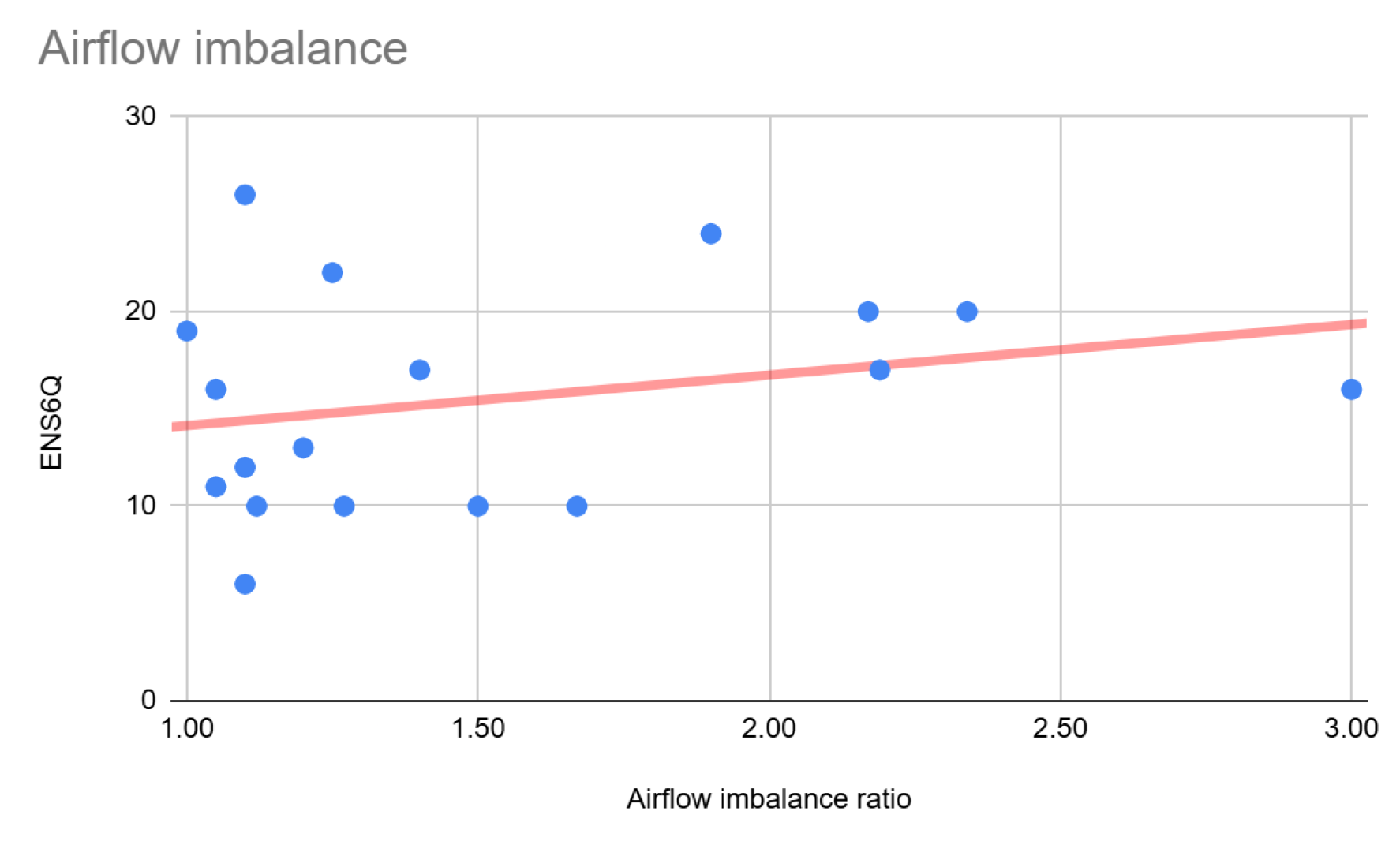

Here, we recorded the airflow on each side of the nasal cavity and calculated a ratio to assess the severity of the imbalance.

Logically, the higher the imbalance, the higher the symptom score, as seen with a weak positive correlation (r = 0.246, p-value = 0.31). High imbalance often represents obstruction on one side and/or a very open side by a very aggressive turbinectomy.

## Discussion

Several interesting things from our point of view emerge from this small study.

First, as several other studies have already pointed out on the ENS, the pressure loss is not well correlated with the severity of the symptoms.(4).

Secondly, the total amount of mucosa strongly correlates with the symptom score. In our opinion, this is an important point raised by this study. Theoretically, a person with more mucosal surface area, thanks to a more developed nasal cavity, will be more resilient in the face of a partial turbinectomy. This could partly explain why with the same amount of cut turbinates some people experience fewer symptoms than others.

Finally, the flow imbalance between each side creates significant discomfort, influencing the symptom severity score.

## Limitations

Our study is limited by the sample size and does not take into account temperature and humidity, which would have been interesting data.

Furthermore, while we have established strong correlations between CFD metrics and symptom severity, direct mucosal nerve density or sensory function measurements were not performed. Future studies incorporating dynamic airflow simulations and biological assessments of mucosal integrity are warranted.

## Conclusion

Our CFD analysis of ENS patients demonstrates that aggressive turbinate reduction leads to a higher ENS6Q score. These findings suggest that preserving mucosal surface area and maintaining symmetrical airflow during nasal surgery may be crucial strategies for minimizing the risk of ENS. Moreover, individuals with a larger preoperative mucosal surface area may be more resilient to turbinectomy-induced changes, experiencing fewer ENS symptoms. Future studies, including advanced simulations and cotton tests, could further refine these strategies and aid in developing targeted interventions for ENS.

